## Supplementary Materials for "Zero-shot learning enables instant denoising and super-resolution in optical fluorescence microscopy"

**This PDF file includes:**

Supplementary Figures 1-15
Supplementary References

### Supplementary Figures

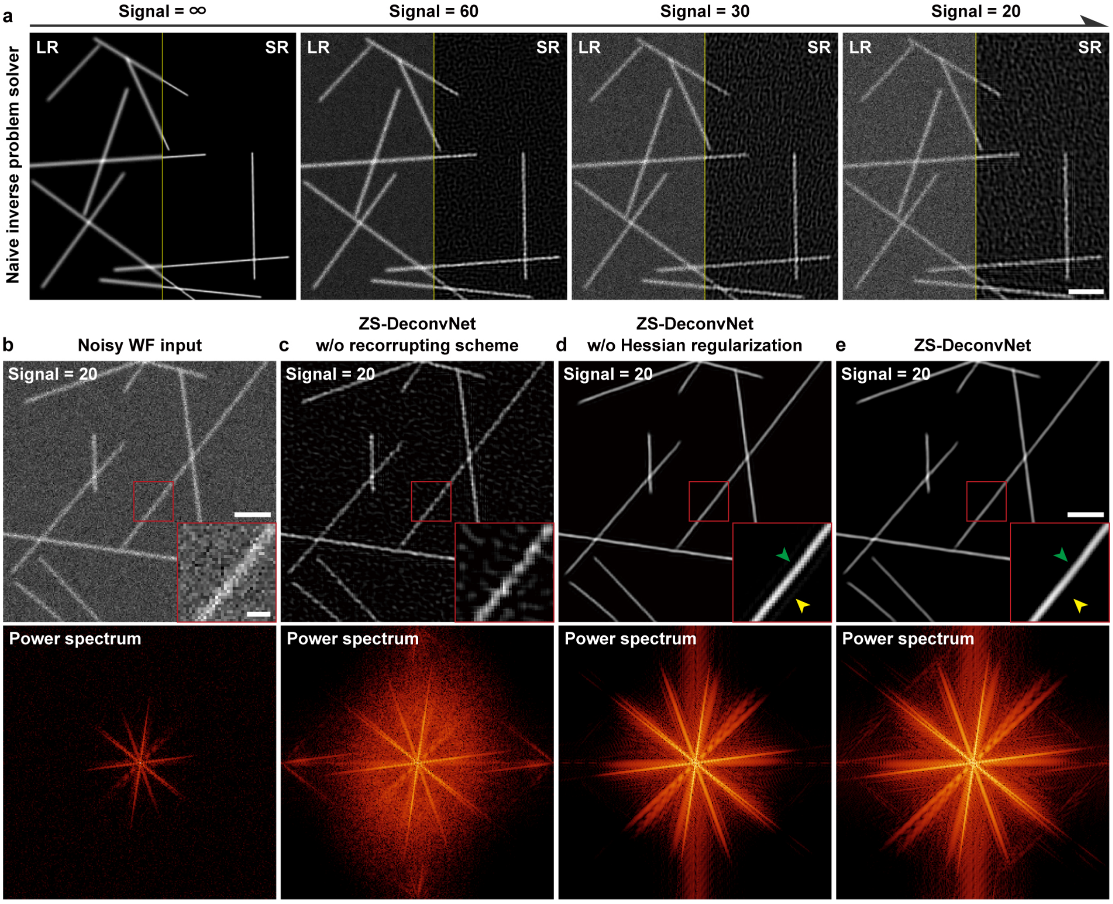


**Supplementary Fig. 1 | Ablation study of ZS-DeconvNet. a**, SR performance for simulated microtubule images with the naïve inverse problem solver under decreasing signal levels of infinite, 60, 30, and 20 (Supplementary Note 3). The displayed images of low resolution (LR, left half) and high resolution (HR, right half) show that the naïve inverse problem solver produces escalating noise-induced artifacts as the signal level decreases. **b-e**, Noisy wide-field image (b) and output images via ZS-DeconvNet (e) and ZS-DeconvNet without the recorrupting scheme (c) and the Hessian regularization (d) under the signal level of 20. The corresponding Fourier power spectrums are shown below, indicating ZS-DeconvNet equipped with both the recorrupting scheme and the Hessian regularization recovers finest HR information. Scale bar, 2 µm (a-e), 0.5 µm (zoom-in regions in b-e).

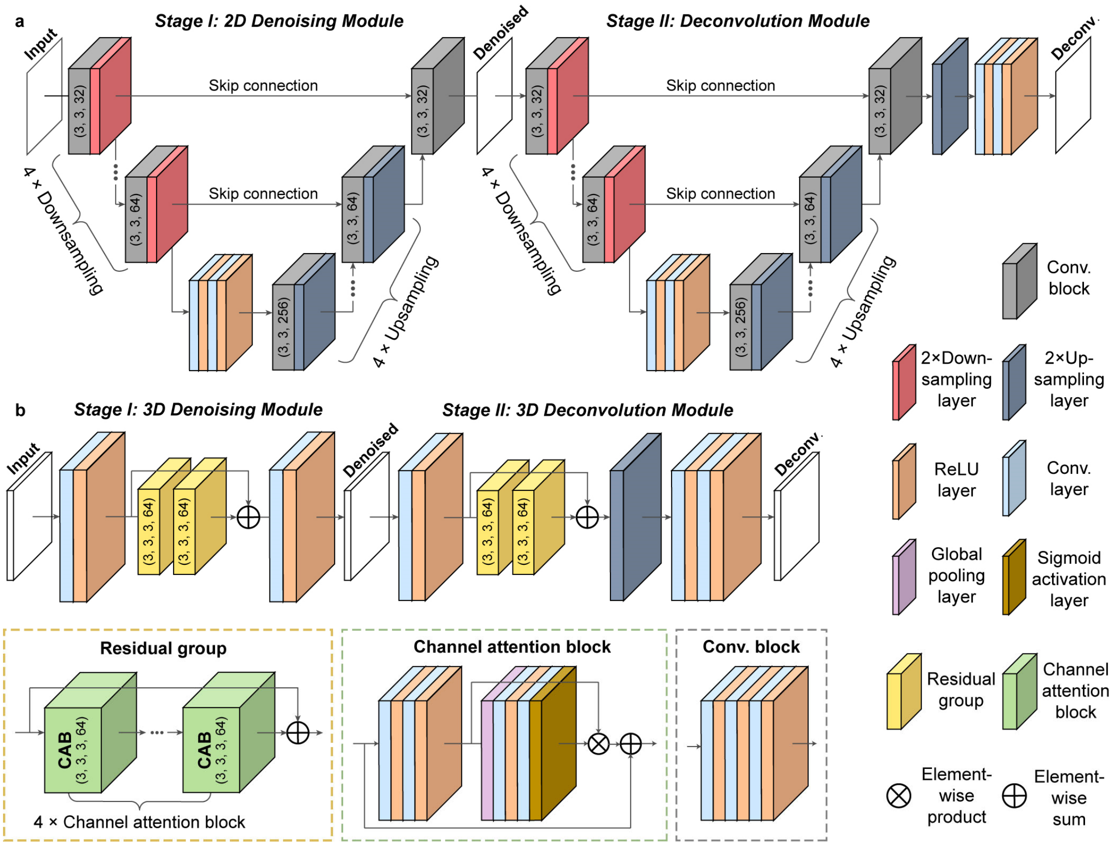


**Supplementary Fig. 2 | Network architectures of ZS-DeconvNet and 3D ZS-DeconvNet. a**, The dual-stage architecture of ZS-DeconvNet used for processing 2D images, which is composed of two sequentially connected U-net models and an up-sampling module including an up-sampling layer and a convolution block. **b**, The dual-stage architecture of 3D ZS-DeconvNet for volumetric data processing. Each stage is composed of a modified 3D residual channel attention network with two residual groups consisting four channel attention blocks. An optional up-sampling module is used to up-sample the feature maps and generate the final monochrome grayscale SR stack.

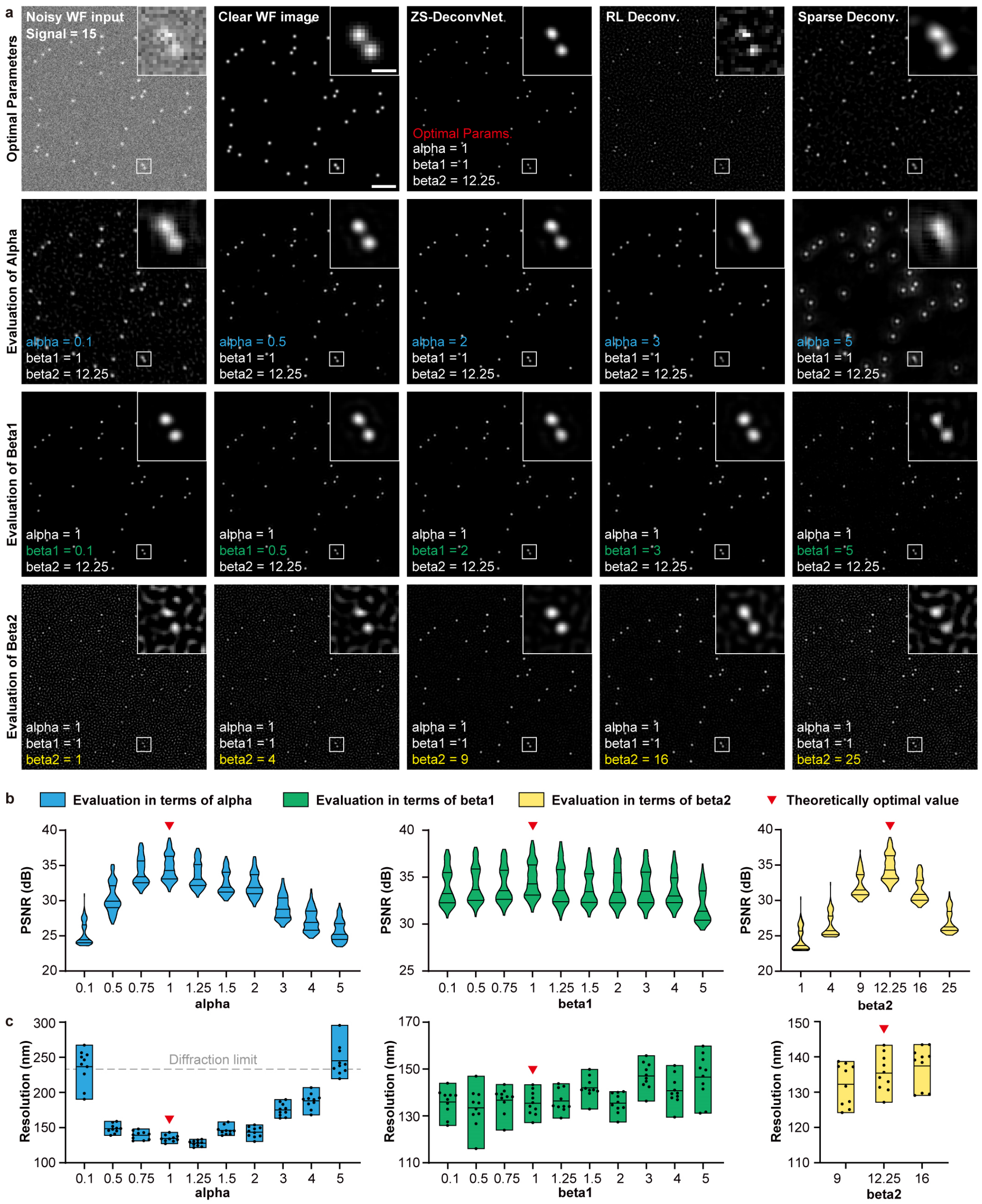


**Supplementary Fig. 3 | Optimal hyperparameter validation for ZS-DeconvNet on simulated images of punctate structures. a**, Images of punctate structures generated by RL deconvolution, sparse deconvolution and ZS-DeconvNet with different hyperparameter choices in terms of alpha (the second row), beta1 (the third row), and beta2 (the fourth row). The noisy wide-field (WF) image was simulated following the steps described in Supplementary Note 2 with a signal level of 15. The diffraction-limited clear image is provided for reference. **b, c**, Statistical evaluations of PSNR (b, n=100) and resolution (c, n=10) for ZS-DeconvNet trained with different hyperparameter choices. The theoretically optimal values for each parameter are labelled with red triangles and the theoretical diffraction limit is labelled with gray dashed lines in c. The resolution was measured with the full width at the half maximum (FWHM) of isolated beads. Both qualitive and quantitative assessments indicate that the experimentally optimal hyperparameters are well consistent with the theoretical analysis in Supplementary Note 1. Scale bar, 2 µm (a), 0.4 µm (zoom-in regions of a).

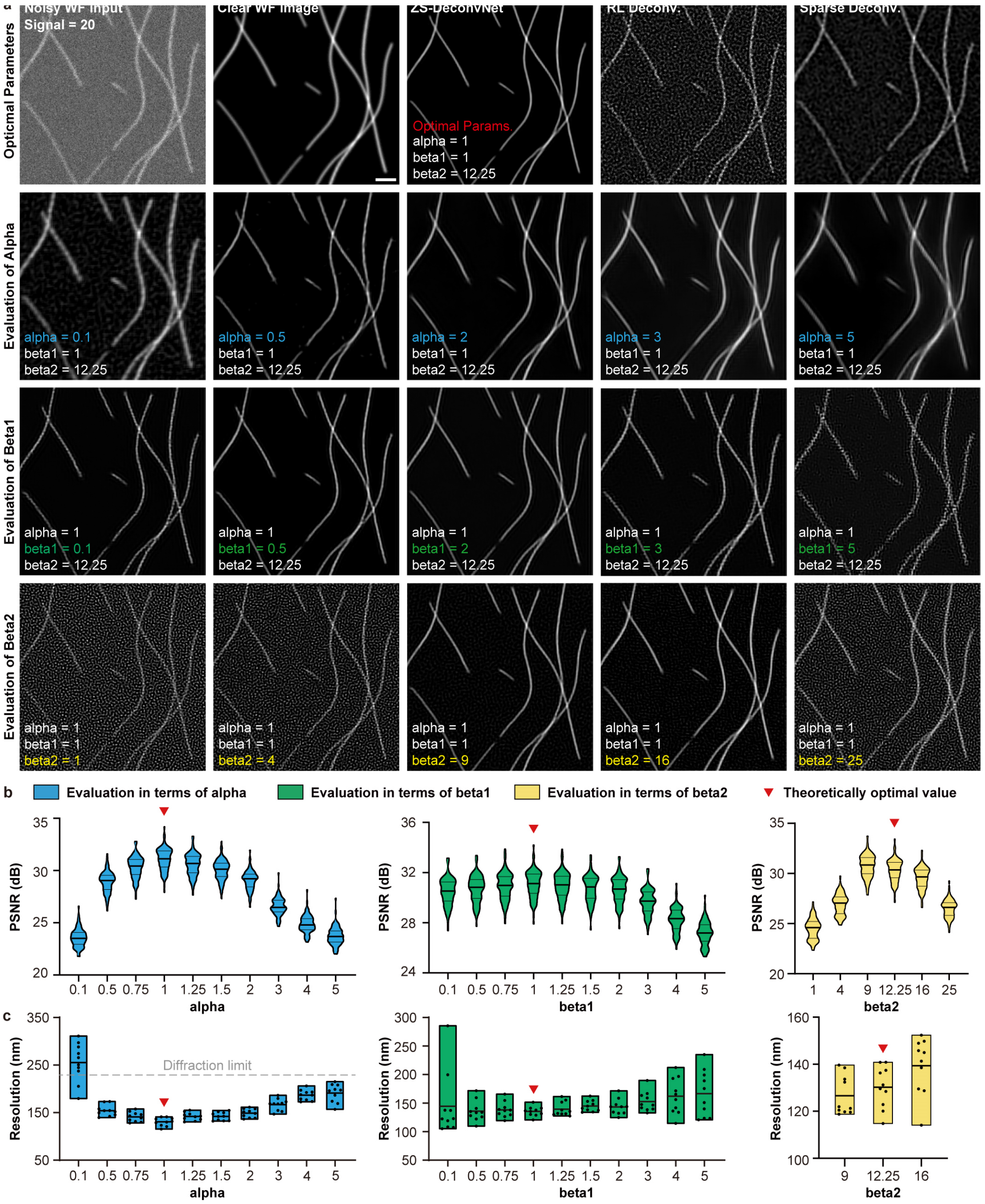


**Supplementary Fig. 4 | Optimal hyperparameter validation for ZS-DeconvNet on simulated images of tubular structures.** **a**, Images of tubular structures generated by RL deconvolution, sparse deconvolution and ZS-DeconvNet with different hyperparameter choices in terms of alpha (the second row), beta1 (the third row), and beta2 (the fourth row). The noisy wide-field (WF) image was simulated following the steps described in Supplementary Note 2 with a signal level of 20. The diffraction-limited clear image is provided for reference. **b, c**, Statistical evaluations of PSNR (b, n=100) and resolution (c, n=10) for ZS-DeconvNet trained with different hyperparameter choices. The theoretically optimal values for each parameter are labelled with red triangles and the theoretical diffraction limit is labelled with gray dashed lines in c. The resolution was evaluated with the full width at the half maximum (FWHM) of isolated microtubules. Both qualitive and quantitative assessments indicate that the experimentally optimal hyperparameters are well consistent with the theoretical analysis in Supplementary Note 1. Scale bar, 1.5 μm.

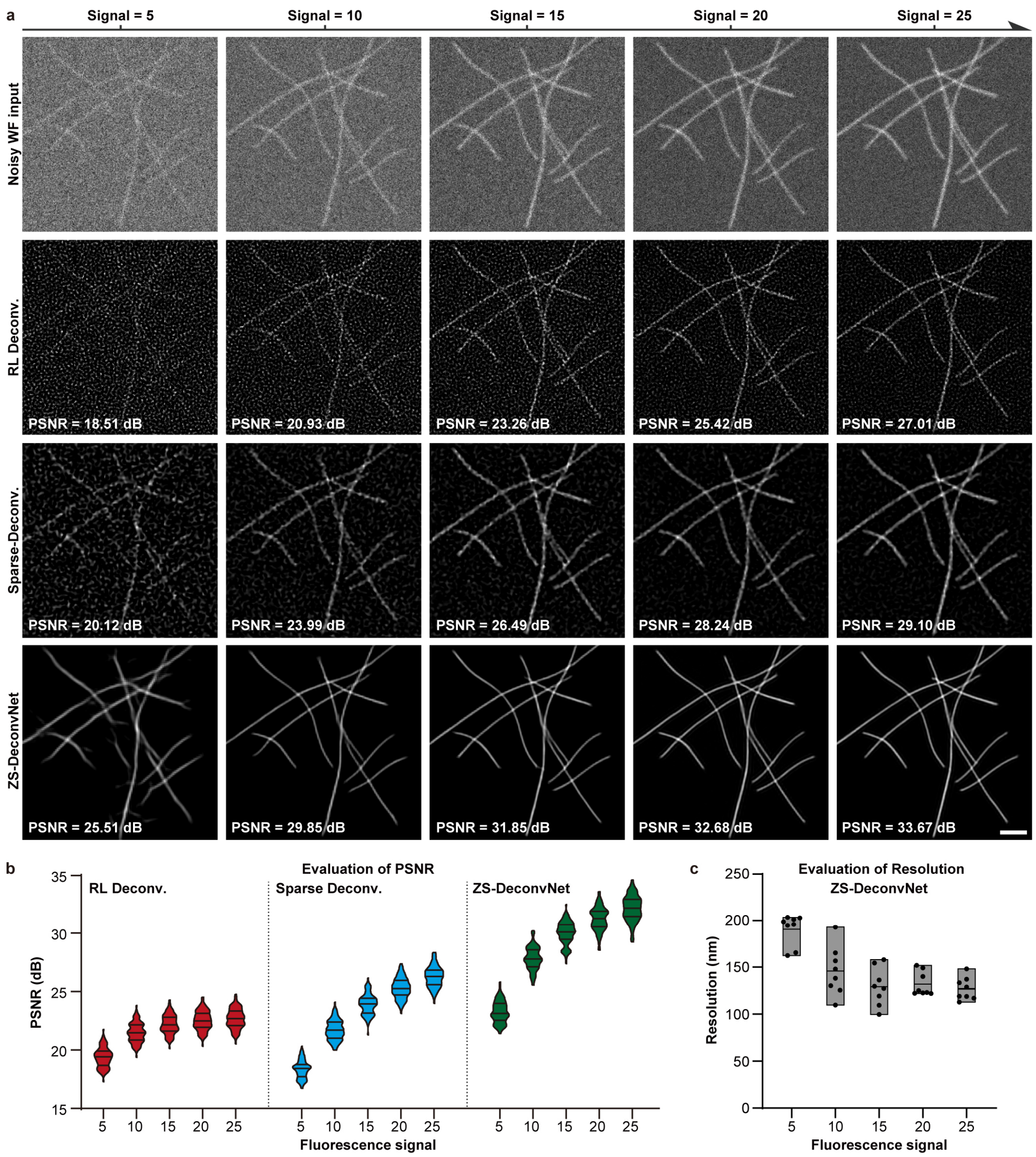


**Supplementary Fig. 5 | Evaluation of ZS-DeconvNet at different fluorescence signal level. a**, Deconvolved images generated via RL deconvolution (the second row), sparse deconvolution (the third row), and ZS-DeconvNet (the fourth row) from noisy wide-filed inputs at fluorescence signal level ranging from 5 to 25 (Supplementary Notes 2). **b**, PSNR comparisons of RL deconvolution (red), sparse deconvolution (blue), and ZS-DeconvNet (green) at different fluorescence signal levels. These results show that ZS-DeconvNet performs state-of-the-art deterministic approaches under various imaging conditions. **c**, Resolution evaluation for ZS-DeconvNet at different fluorescence signal levels, which were measured with the FWHM of sectioned profiles. Scale bar, 2 μm.

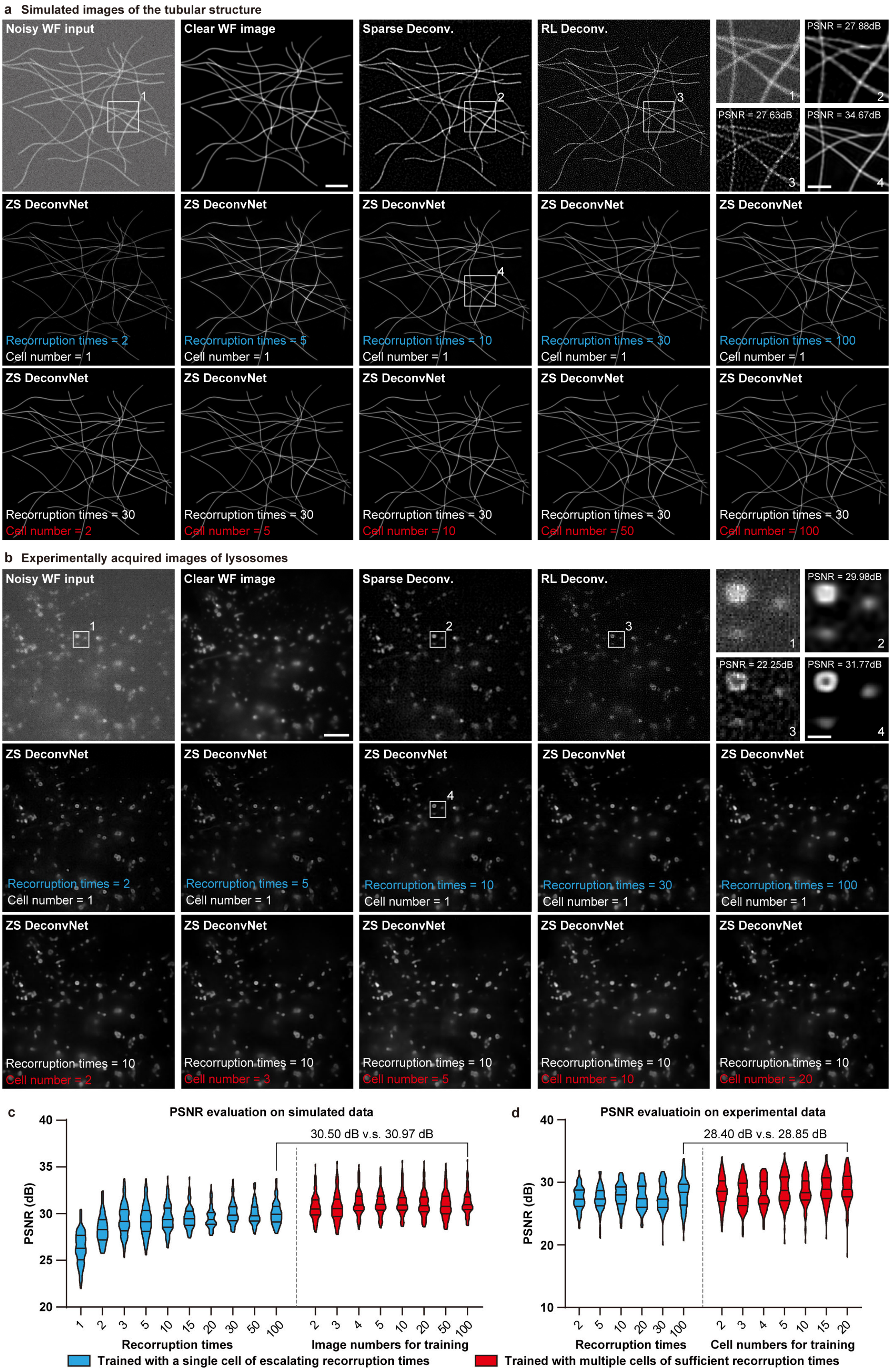


**Supplementary Fig. 6 | Evaluation and characterization of ZS-DeconvNet trained with different augmentation strategies and scales of dataset. a,** Deconvolved images of simulated tubular structures enhanced by RL deconvolution, sparse deconvolution, and ZS-DeconvNet trained on a single image with different recorruption times (ranging from 2 to 100) and different amounts of images (ranging from 2 to 100) with a fixed recorruption time of 30. The diffraction limited clear image is shown for comparison. **b**, Deconvolved images of lysosomes enhanced by RL deconvolution, sparse deconvolution, and ZS-DeconvNet trained on a single image with different recorruption times (ranging from 2 to 100) and different amounts of images (ranging from 1 to 20) with a fixed recorruption time of 10. The diffraction limited clear image is shown for comparison. **c**, **d**, Statistical evaluation of ZS-DeconvNet on simulated data (c, n=100) and experimental data (d, n=100) in terms of PSNR trained with different recorruption times (blue) and escalating scales of dataset (red). Of note, the ZS-DeconvNet models trained with a single image recorrupted with a sufficient time show comparable performance to the models trained with abundant training dataset for both simulated and experimental data, i.e., decreasing less than 0.5 dB in PNSR, indicating the capability and robustness of ZS-DeconvNet even trained with a single input image. Scale bar, 4 μm (a, b), 1.5 μm (zoom-in region in a), 0.4 μm (zoom-in region in b).

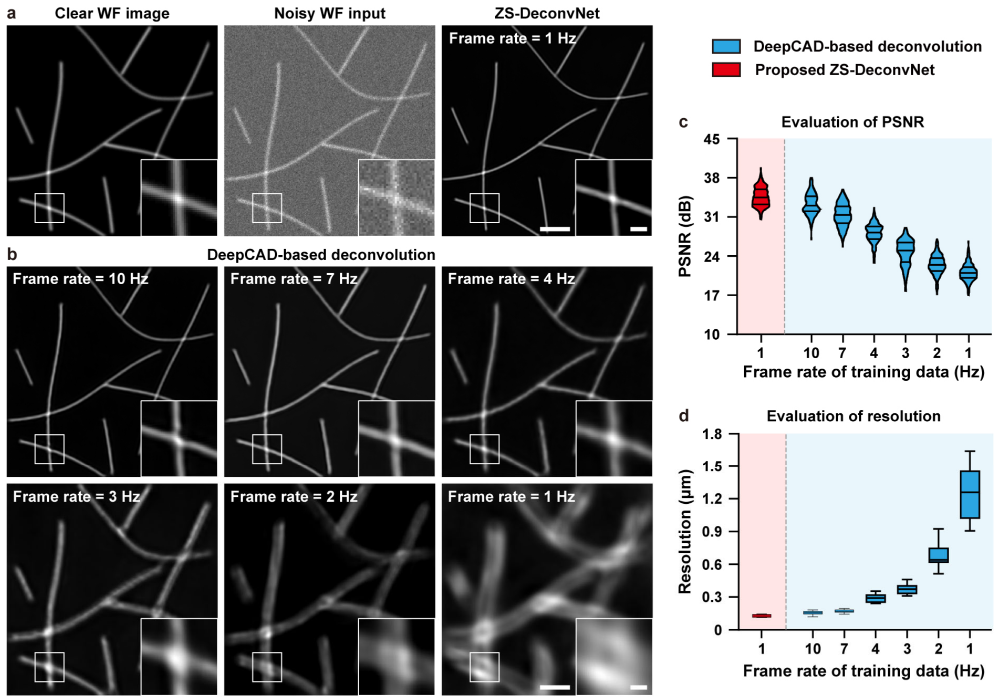


**Supplementary Fig. 7 | Comparison of ZS-DeconvNet and DeepCAD-based deconvolution networks on simulated time-lapse images of tubular structures. a**, Representative wide-field frame with/without noises and the corresponding ZS-DeconvNet enhanced image from a simulated 1000-frame video of tubular structures at a 1Hz frame rate. The average moving speed of simulated microtubules is set to ~1 µm/s (Supplementary Note 2), which is approximately consistent with the growth velocity of microtubules in live COS-7 cells[1]. The ZS-DeconvNet model was trained with all frames of the video. **b**, Deconvolved outputs of the same image shown in a, which were produced with DeepCAD-based deconvolution networks (Methods) trained on simulated time-lapse data of different imaging frame rate ranging from 1 Hz to 10 Hz. **c**, **d**, Statistical comparison of ZS-DeconvNet used in a (red) and DeepCAD-based deconvolution networks used in b (blue) in terms of (c) PSNR (n=100) and (d) resolution (n=15). The resolution was evaluated with the FWHM of microtubules and the theoretical diffraction limit is labelled with gray dashed lines in d. These results together suggest that ZS-DeconvNet can be well-trained with time-lapse data lacking of temporal continuity, yielding superior inference fidelity and resolution compared with recently proposed self-supervised learning schemes for microscopy data, i.e., DeepCAD. Scale bar, 2 µm (a, b), 0.5 µm (zoom-in regions of a and b).

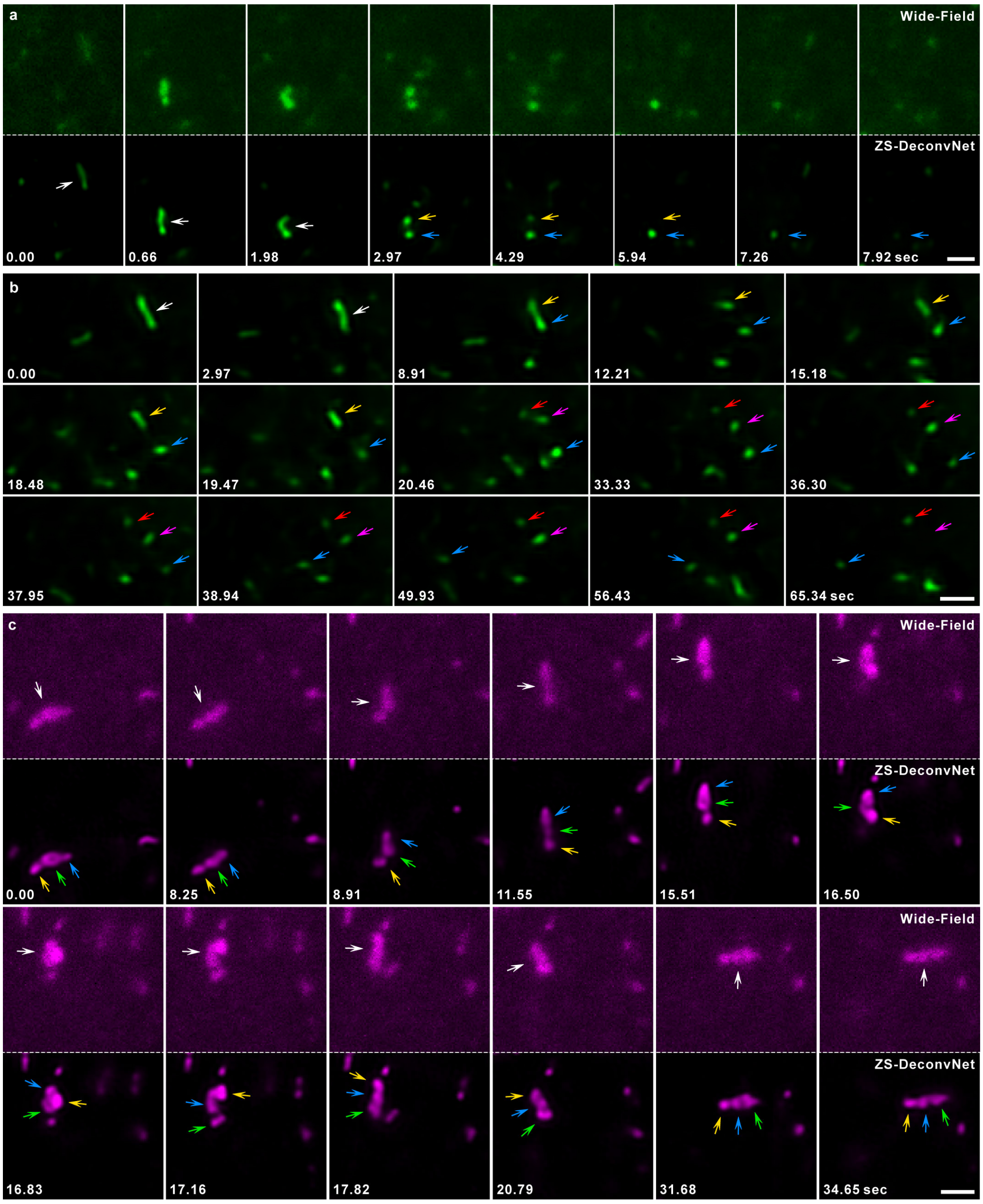


**Supplementary Fig. 8 | Interesting showcases of the dynamics of recycling endosomes (REs) and lysosomes or late-endosomes (Lyso/LEs) in live SUM159 cells revealed by ZS-DeconvNet. a**, Time-lapse images of a fission event of RE, and both of the divided REs undergo exocytosis sequentially (indicated by yellow and blue arrows, respectively). The corresponding noisy wide-field images are provided in the upper row for comparison. **b**, Time-lapse images of another fission event of RE, in which a tubular RE divides into three parts and one of them (indicated by blue arrows) ran away independently. **c**, Time-lapse images of three tethered Lyso/LEs traveling together, and rearranging their sequence in the period of 15.51 to 17.82 sec. The corresponding noisy wide-field images are shown in the upper rows, where the tethering details and the rearrangement of the three Lyso/LEs can hardly be recognized. Scale bar, 1 μm (a, b), 1.5 μm (c).

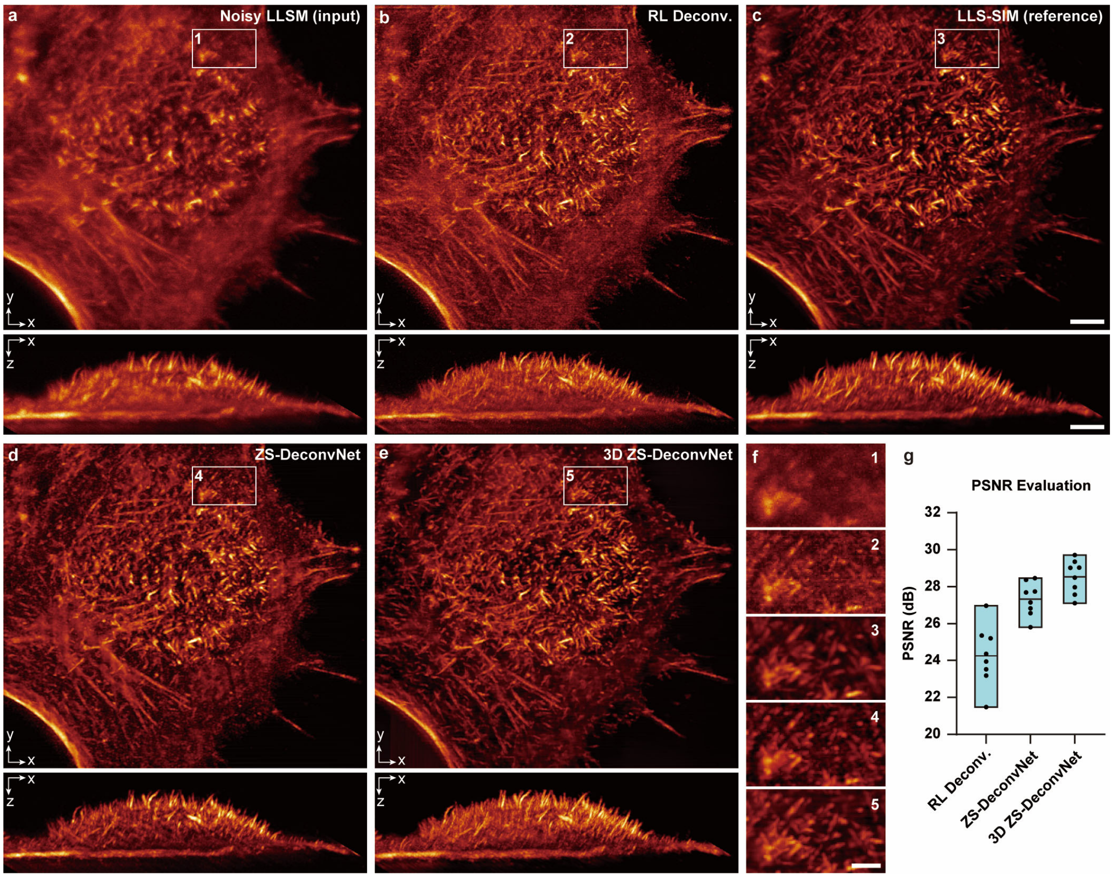


**Supplementary Fig. 9 | Comparison of SR capability for 3D LLSM images with RL deconvolution, ZS-DeconvNet, and 3D ZS-DeconvNet. a-e**, Representative maximum intensity projections (MIP) of F-actin in a COS-7 cell imaged by (a) LLSM (low excitation), (c) LLS-SIM (high excitation) and reconstructed with (b) RL deconvolution, (d) recorruption-based ZS-DeconvNet, and (e) spatially interleaved self-supervised 3D ZS-DeconvNet from the noisy LLSM image stack. Both *xy*-MIPs (upper) and *xz*-MIPs (lower) are provided. **f**, Magnified regions labelled in a-c with white boxes. g, Statistical comparison in terms of PSNR for RL deconvolution, ZS-DeconvNet, and 3D-DeconvNet (n=8). Scale bar, 8 μm (a-e), 2 μm (f).

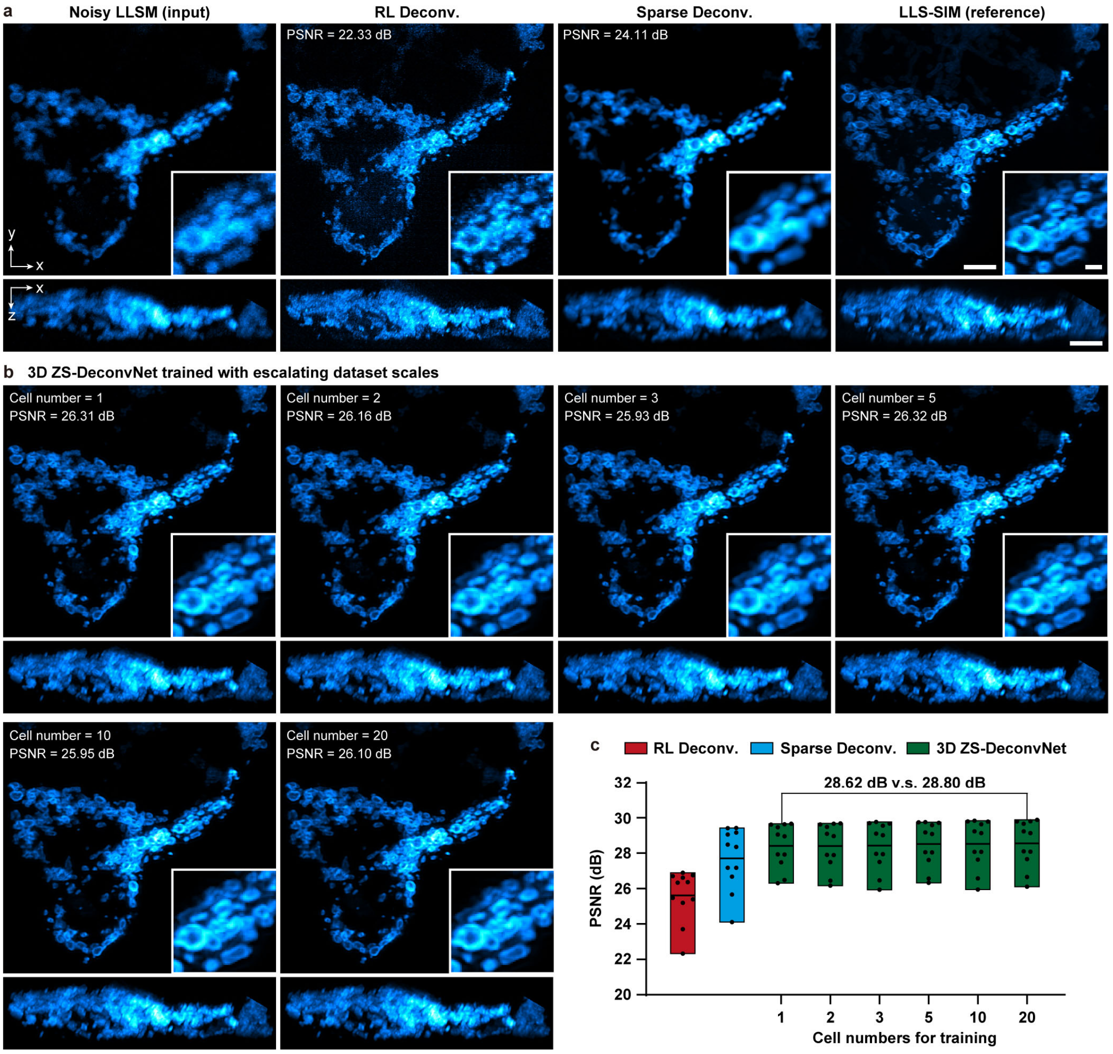


**Supplementary Fig. 10 | Evaluation of 3D ZS-DeconvNet trained with different scales of datasets.** **a**, Representative maximum intensity projections (MIP) of mitochondrial outer membrane in a 293T cell imaged by LLSM (low excitation), LLS-SIM (high excitation) and reconstructed with RL deconvolution, sparse deconvolution from the noisy LLSM image stack. **b**, Super-resolved MIP images generated by 3D ZS-DeconvNet models trained with escalating dataset scales of a single cell to 20 cells. **c**, Statistical comparisons in terms of PSNR for RL deconvolution (red), sparse deconvolution (blue), and 3D ZS-DeconvNet trained with different dataset scales (green). Both *xy*-MIPs (upper) and *xz*-MIPs (lower) are shown and the PSNR values are labelled in the top left corner of each image. These results suggest the successful zero-shot implementation of 3D ZS-DeconvNet where the model was trained with data augmented from a single image stack, while yielding comparable performance to models trained with more data and outperforming conventional deconvolution algorithms. Scale bar, 5 µm (*xy*- and *xz*-MIPs of a and b), 1 µm (zoom-in regions of a and b).

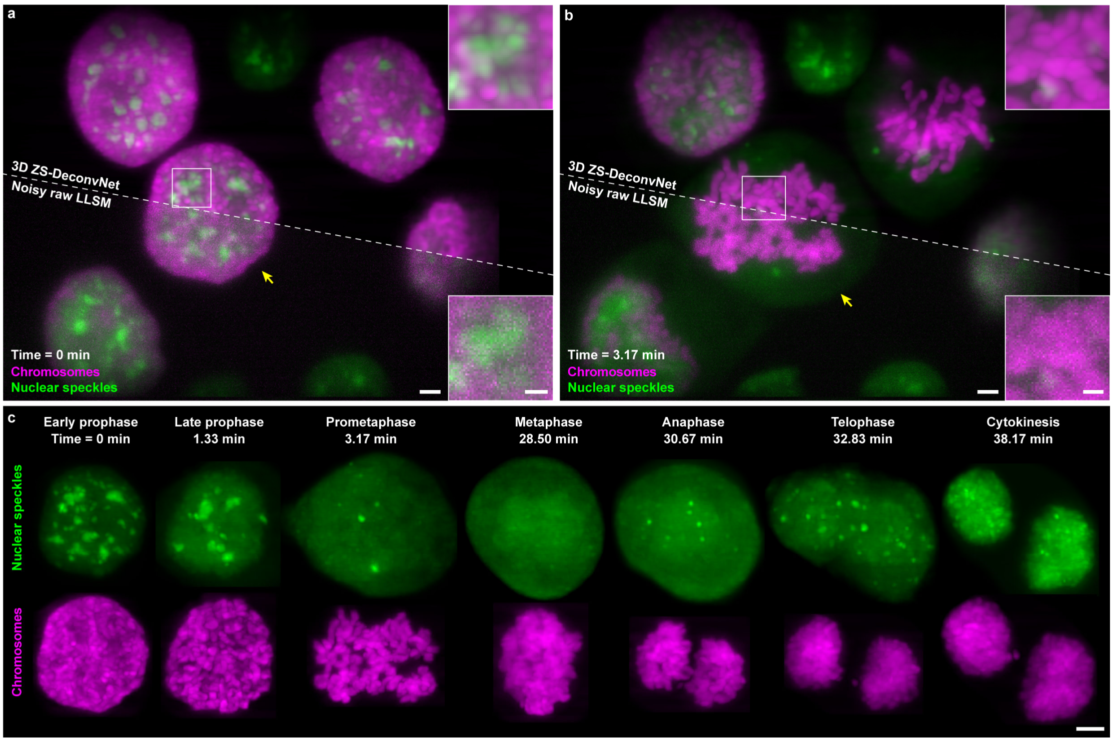


**Supplementary Fig. 11 | Visualizing the behaviors of HeLa-SC35 labelled nuclear speckles and mCherry-H2B labelled chromosomes during cell mitosis via 3D ZS-DeconvNet enhanced LLSM.** **a**, **b**, Two representative frames imaged by LLSM with low light-dose (bottom left) and enhanced by 3D ZS-DeconvNet (top right) in early prophase and prometaphase, respectively, during mitosis of several HeLa cells. **c**, Time-lapse 3D ZS-DeconvNet enhanced images of a mitotic HeLa cell indicated with yellow arrow in a and b, labelled by HeLa-SC35 and mCherry-H2B, showing the disassemble and reassemble process of nuclear speckles during mitosis (Supplementary Video 5). Scale bar, 2.5 μm (a and b), 1 μm (zoom-in regions of a and b), 4 μm (c).

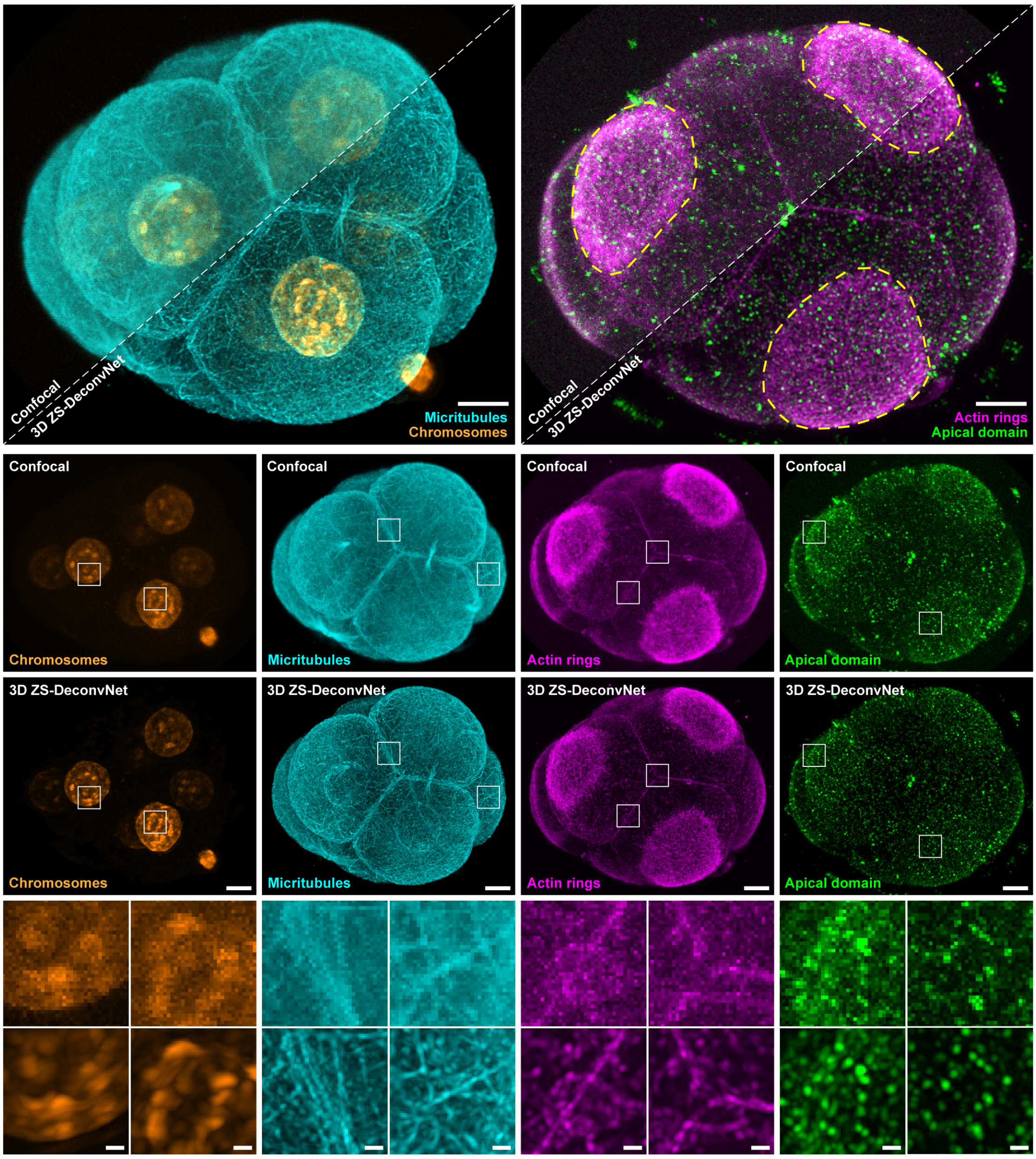


**Supplementary Fig. 12 | Four-color super-resolution visualization of another early mouse embryo via 3D ZS-DeconvNet.** 3D-rendering confocal images of early mouse embryo immunostained for microtubule bridges (cyan), chromosomes (orange), actin rings (magenta), and apical domain (green) before and after 3D ZS-DeconvNet enhancement. The 3D ZS-DeconvNet models trained with the input noisy data itself provide a dramatic improvement in both SNR, contrast, and resolution compared to the original confocal image stack. Scale bar, 8 µm (images of the entire embryo), 1 µm (zoom-in regions for each color channel).

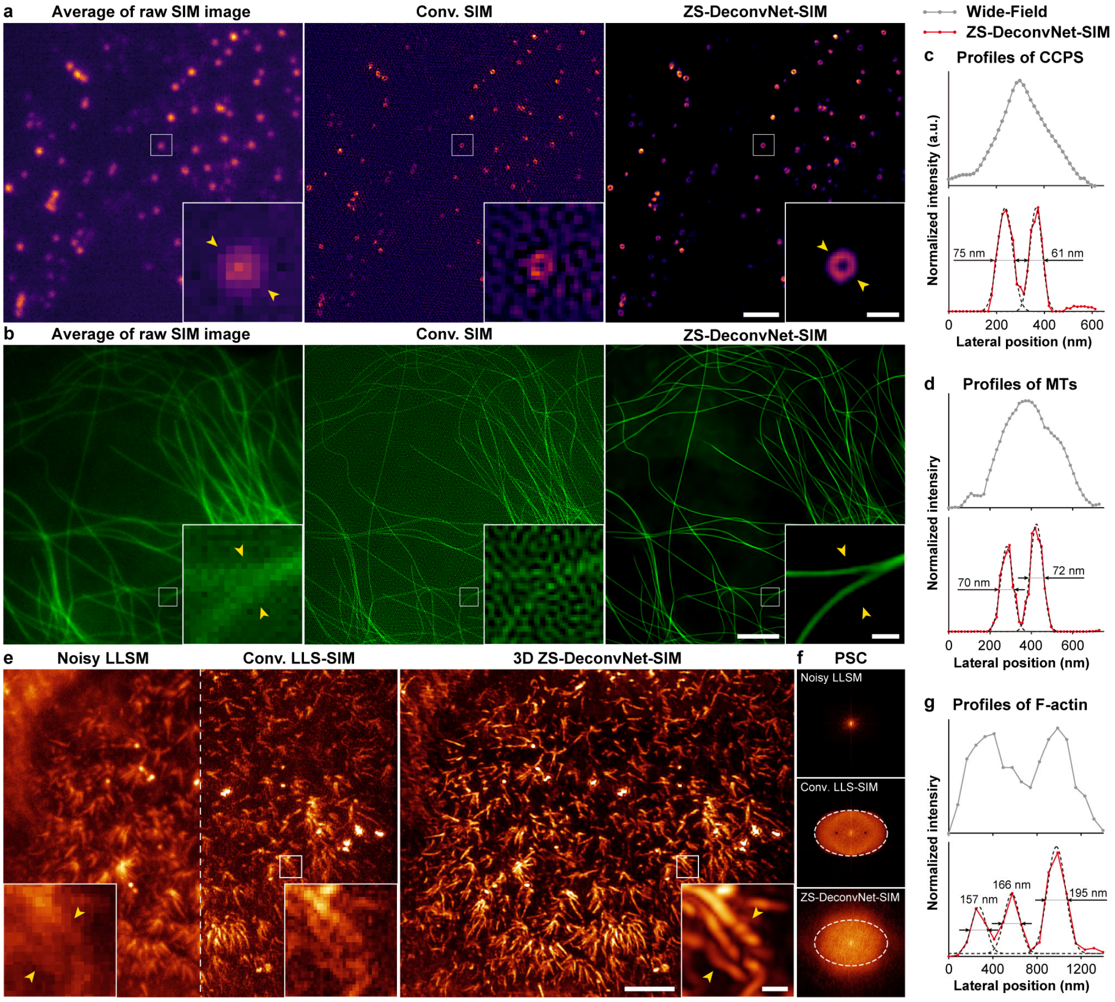


**Supplementary Fig. 13 | Resolution comparison between wide-field images, conventional SIM images, and ZS-DeconvNet enhanced SIM images across multiple SIM modalities. a, b**, Representative SR images of (a) clathrin coated pits (CCPs) and (b) microtubules (MTs) acquired with the (a) TIRF-SIM and (b) GI-SIM mode, respectively, of our Multi-SIM system and reconstructed with the conventional SIM algorithm and ZS-DeconvNet. **e, f**, Representative (e) SR images and (f) corresponding power spectrum coverages (PSC) of F-actin acquired with the LLS-SIM system and reconstructed via the conventional LLS-SIM algorithm and 3D ZS-DeconvNet. The envelopes of the LLS-SIM image are labelled with white dashed ellipse in d, suggesting an isotropic super-resolution capability of 3D-DeconvNet-SIM. **c, d, g**, Intensity profile plots for wide-field images (gray, upper panels) and ZS-DeconvNet-SIM images (red, lower panels) of (c) CCPs, (d) MTs, and (g) F-actin along the lines indicated by the two yellow arrowheads in a–c. In these cases, the hollow structures of CCPs and adjacent filaments of microtubules and F-actin were clearly resolved by ZS-DeconvNet-SIM with a spatial resolution of ~70 nm for 2D-SIM and ~160 nm for LLS-SIM. By contrast, they were distinguishable with either LLSM for its resolution limitation or conventional LLS-SIM for artifact contamination. Scale bar, 3 μm (a), 0.5 μm (zoom-in region of a), 3 μm (b), 0.3 μm (zoom-in region of b), 5 μm (e), 1 μm (zoom-in region of e).

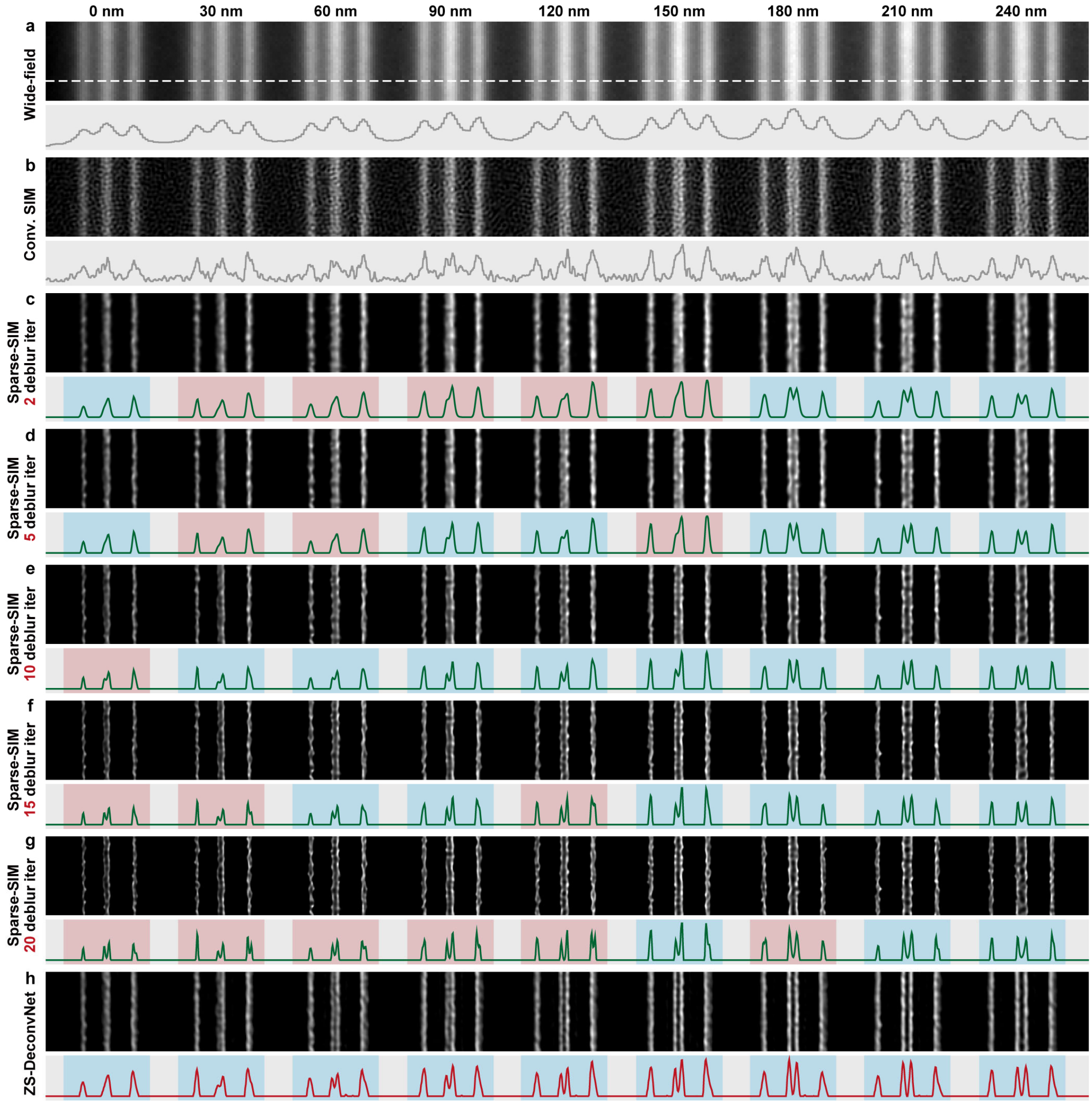


**Supplementary Fig. 14 | Comparison between ZS-DeconvNet-SIM and sparse-SIM on Argo-SIM slides. a**, **b**, Wide-filed (a) and convolutional SIM (Conv. SIM) images (b) of Argo-SIM slides which consists 9 pairs of dual lines, whose spacing gradually increases from 0 nm to 240 nm with a step of 30 nm. **c-g**, Sparse-SIM images reconstructed from the Conv. SIM image shown in b with different deblur iterations of ranging from 2 to 20. **h**, SR image of the same content generated via ZS-DeconvNet-SIM model trained with noisy data only. The intensity profiles along the white dashed line labeled in a are shown in the lower panel for each method. We highlighted correct and false reconstructions including failing to resolve two parallel lines or wrongly distinguishing a single line apart, with bottom colors of blue and red on the line profiles of sparse-SIM and ZS-DeconvNet-SIM images, respectively. Of note, with a relatively large deblur iteration, i.e., larger than 10 (e-g), sparse-SIM generates several false-positive errors where a single line was wrongly resolved into two separated lines, while the proposed ZS-DeconvNet-SIM does not require any user-defined parameters and thereby yields SR images with higher fidelity.

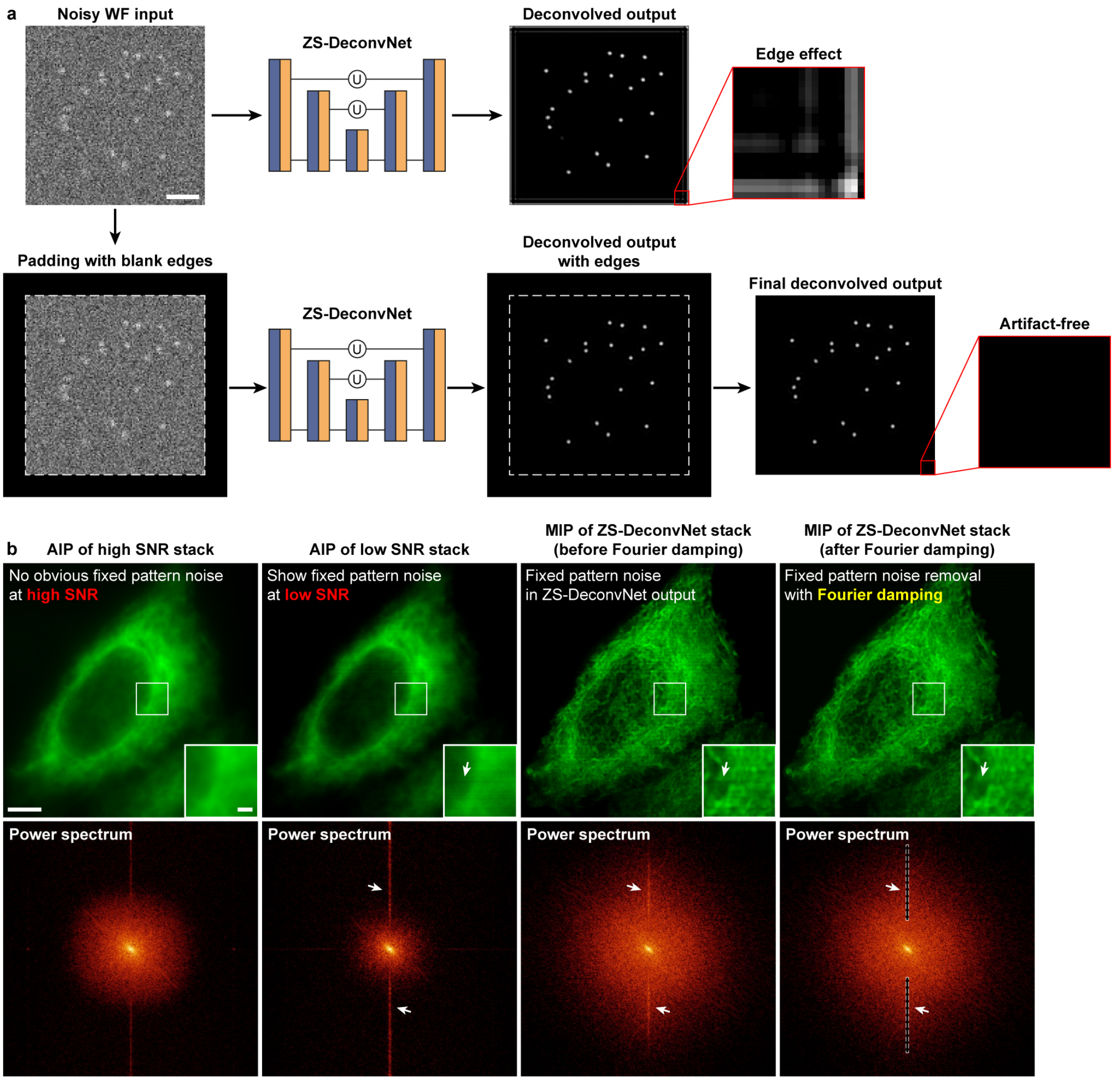


**Supplementary Fig. 15 | Artifact elimination for ZS-DeconvNet. a**, A representative case showing that the deconvolution-induced edge artifact (upper row) and the artifact elimination by padding blank edges surrounding the input image and cutting them off after network processing (lower row). **b**, Representative average intensity projections (AIPs) of ER acquired by LLSM at high (first column) and low (second column) SNR conditions, and corresponding MIPs of 3D ZS-DeconvNet enhanced image stacks before (third column) and after (fourth column) the Fourier damping operation. The power spectrums of each image are shown below. The emergence and removal of fixed pattern noises is highlighted by white arrows in spatial and Fourier domains and the Fourier apodization masks are labelled with white dashed rectangles. Scale bar, 1.5 μm (a), 5 μm (b), 1 μm (zoom-in region of b).

#### Supplementary Tables

##### Supplementary Table 1. Implementation details of ZS-DeconvNet

| | Imaging method | Network model type | Initial learning rate | Training patch size | Training batch size | Total training iterations | Training time (hours) |
|---|---|---|---|---|---|---|---|
| Fig. 1c | TIRF | ZS-DeconvNet | $5 \times 10^{-5}$ | $144 \times 144$ | 4 | 80,000 | 3 |
| Fig. 2a-d<br>Supplementary Videos 2, 3 | TIRF | ZS-DeconvNet | $5 \times 10^{-5}$ | $128 \times 128$ | 4 | 50,000 | 1 |
| Fig. 2e, f, i<br>Supplementary Fig. 8<br>Supplementary Video 4 | TIRF | ZS-DeconvNet | $5 \times 10^{-5}$ | $144 \times 144$ | 4 | 50,000 | 2 |
| Fig. 3c | LLSM | 3D ZS-DeconvNet | $1 \times 10^{-4}$ | $64 \times 64 \times 13$ | 3 | 10,000 | 3.5 |
| Fig. 3e, f<br>Supplementary Video 5 | LLSM | 3D ZS-DeconvNet | $1 \times 10^{-4}$ | $64 \times 64 \times 13$ | 3 | 10,000 | 3.5 |
| Fig. 4a-d<br>Supplementary Fig. 12<br>Supplementary Video 7 | Confocal microscopy | 3D ZS-DeconvNet | $1 \times 10^{-4}$ | $48 \times 48 \times 5$ | 3 | 10,000 | 2 |
| Fig. 4e-h<br>Supplementary Video 8 | 3D wide-field microscopy | 3D ZS-DeconvNet | $1 \times 10^{-4}$ | $64 \times 64 \times 13$ | 3 | 10,000 | 4 |
| Fig. 5b, c<br>Supplementary Fig. 13a, b | TIRF-SIM<br>GI-SIM | ZS-DeconvNet-SIM | $5 \times 10^{-5}$ | $128 \times 128$ | 4 | 50,000 | 1 |
| Fig. 5d, e<br>Supplementary Fig. 13c | LLS-SIM | 3D ZS-DeconvNet-SIM | $1 \times 10^{-4}$ | $64 \times 64 \times 13$ | 3 | 10,000 | 4 |
| Supplementary Fig. 10 | LLSM | 3D ZS-DeconvNet | $1 \times 10^{-4}$ | $64 \times 64 \times 13$ | 3 | 10,000 | 3.5 |
| Supplementary Fig. 11<br>Supplementary Video 6 | LLSM | 3D ZS-DeconvNet | $1 \times 10^{-4}$ | $64 \times 64 \times 13$ | 3 | 10,000 | 3.5 |

#### Supplementary Table 2. Imaging conditions of live-cell experiments

| | Imaging method | Sample | Label | Excitation NA | Excitation λ (nm) | Exposure time per raw image (ms) | Total acquisition time | Illumination intensity | Cycle time (Acquisition + resting time) (sec) | Time points |
|---|---|---|---|---|---|---|---|---|---|---|
| Figs. 2a, b Supplementary Video 2 | TIRF microscopy | COS-7 | lifeact-mEmerald myosion2-Halo-JF549 | 1.41 | 488, 560 | 5 | 0.19s | 488: 2 $W/cm^2$<br>560: 10.8 $W/cm^2$ | 5 | 110 |
| Figs. 2c, d Supplementary Video 3 | TIRF microscopy | COS-7 | lifeact-mEmerald myosion2-Halo-JF549 | 1.41 | 488, 560 | 5 | 0.19s | 488: 1.7 $W/cm^2$<br>560: 10.8 $W/cm^2$ | 5 | 825 |
| Fig. 2e, f, i Supplementary Video 4 | TIRF microscopy | SUM-159 | EGFP-Rab11 Lamp1-Halo-JF549 | 1.35 | 488, 560 | 1 | 0.086s | 488: 119-412 $W/cm^2$<br>560: 57.1-149 $W/cm^2$ | 0.33 | 1499 |
| Fig. 3e, f Supplementary Video 5 | LLSM | HeLa | Calnexin-mEmerald H2B-Halo-JF642 Mito-dsRed | 0.25, 0.14 | 488, 560, 642 | 10 | 5.904s | 488: 1.39 μW<br>560: 0.63 μW<br>642: 0.14 μW | 10 | 937 |
| Fig. 4e-h Supplementary Video 8 | 3D WF microscopy | C. elegans embryo | wyEx51119, jcIs1 qxIs257 | 1.35 | 488, 560 | 10 | 8.705s | 488: 2.4 mW<br>560: 3.6 mW | 30 | 213 |
| Supplementary Fig. 11 Supplementary Video 6 | LLSM | HeLa | SC35-mEmerald H2B-mCherry | 0.25, 0.14 | 488, 560 | 10 | 3.706s | 488: 2.46 μW<br>560: 9.29 μW | 30 | 318 |
